# RecNW: A fast pairwise aligner for targeted sequencing

**DOI:** 10.1101/371989

**Authors:** Alexandre Yahi, Tuuli Lappalainen, Pejman Mohammadi, Nicholas P. Tatonetti

## Abstract

**Motivation:** Targeted sequencing aims at in-depth analysis of specific genomic loci through high-throughput sequencing for applications such as resequencing or CRISPR gene editing. These applications require exact pairwise alignment algorithms to fully characterize large amounts of reads by comparison to the targeted locus, or reference. Optimal solutions to this alignment problem are provided by classic implementations of the global and semi-global versions of Needleman-Wunsch algorithms, but they remain computationally expensive due to their quadratic complexity in time and space.

**Implementation:** In this paper we present RecNW, an open source C++ exact aligner packaged for Python that implements the semi-global version of the Needleman-Wunsch algorithm with affine gap penalty. RecNW utilizes low complexity of targeted sequencing libraries by aligning only unique reads, and recurrently using blocs of the alignment matrix between reads based on their similarities. Through this, RecNW performs exact alignment on average more than four times faster than gold standard comparable software.

**Software:** https://github.com/AYahi/recNW

## 1 Introduction

Pairwise alignment aims at comparing two sequences and finding their similarities. Symbols of each sequence can either be aligned to represent a match or mismatch, or a gap can be inserted to represent an insertion or a deletion. Dynamic programming is used for calculating optimal sequence alignment efficiently. There are two variations of this approach that are widely used: the Needleman-Wunsch global pairwise aligner [11] that aligns sequences end-to-end and Smith-Waterman local algorithm[12] that tries to match a sequence to a substring of the reference sequence it is compared to. Other methods such as Basic Local Alignment Search Tool (BLAST) [1] or Burrows-Wheeler Aligner (BWA) [8] use heuristic approaches to the pairwise alignment problem, and they offer in general superior time performances necessary for genome-wide alignment.

However, many study designs use targeted sequencing, where a locus of interest is captured or typically amplified by PCR, and sequenced on a high-throughput sequencing platform such as Illumina MiSeq or HiSeq. This approach is used in applications such as targeted panel sequencing for mutation detection [10], integrative taxonomy [3], as well as targeted genome editing by CRISPR and other technologies [4, 2, 6]. The data of targeted sequencing is often comprised of very high numbers of reads from a relatively small known locus. For the alignment of such read data to the known locus of origin, exact methods are preferred for precision at calling indels and mutations [7, 13], in spite of their comparatively lower time performance against heuristic methods.

In this paper we present RecNW (Recursive Needleman-Wunsch), an exact pairwise aligner based on the semi-global variation of Needleman-Wunsch algorithm that supports affine gap penalty [5]. RecNW utilizes the high level of read similarity between reads that is inherent to targeted sequencing libraries to speed up the alignment process. We benchmark performance of RecNW against gold standard tools under a range of scenarios using extensive simulated libraries and real world data from polyclonal and monoclonal CRISPR gene editing experiments. RecNW is implemented in C++ and packaged for Python, and distributed under the Apache License 2.0 license.

## 2 Methods

RecNW is an implementation of Needleman-Wunsch algorithm, with the Gotoh variation for affine gap penalty that makes the distinction between gap opening and gap extension to encourage less gaps of higher length rather than many short gaps.

The input data to RecNW is a sequencing read file in fastq format, and the reference sequence for the targeted locus. The input reads are assumed to be trimmed for sequencing adaptors and low-quality bases if needed. Unique reads are identified and sorted using Python’s build-in sort function. This step relies on the Timsort algorithm, a hybrid of merge sort and insertion sort algorithm. Semi-global alignment strategy was adopted to achieve compact alignment of entire short reads to their corresponding position in the relatively larger reference locus. This was implemented by initializing the alignment matrices with zeros on the first row and first column for no head gap penalty, and starting the traceback to the highest score of the last row or column for free tail gaps.

To further utilize the inherent redundancy in the read library, RecNW was implemented to reuse a part of the alignment matrix from each read for the next read. Specifically, in a sorted list of unique reads, the *N* first rows of the alignment matrix for the ith read are carried over to the matrix for the *i* + 1^*th*^ read, with N being the length of the longest nucleotide shared head sequence between reads *i* and *i* + 1 while the reference sequence remains identical. This leverages on the similarity between reads to reduce the computation time.

We evaluated RecNW against NEEDLE by first aligning 3.7M reads across eight datasets (with 292,128-632,834 reads) of polyclonal and monoclonal CRISPR gene editing experiments on the FLCN gene (chr17: 17214783-17215323 in hg38) from Illumina MiSeq. We then generated 640 synthetic datasets of 150 bps sequences, using a locus of the ACCS gene (chr11: 44086824-44087153 in hg38) for an in depth evaluation of the alignment speed. We generated these datasets by changing the number of reads (5k, 50k, 500k, 5M), the average number of indels (1, 1.5, 2, 3 per read), and the range of position where indels are allowed on the sequence (from 0, 25, 50 and 75 base pairs to the end). These different input parameters allowed for a variety of realistic percentages of unique reads and percentages of alignment matrix lines reused based on sample-to-sample head similarity. We ran all experiments on a server with 64 CPU cores AMD Opteron 6272 and 322 Gb of RAM running under Ubuntu 14.04 with Python 2.7.8, g++ 4.8.4, Cython 0.24, NumPy 1.11.3 and SciPy 0.19.1.

## 3 Results

We compared the output from our RecNW aligner that reuses lines of the alignment matrix between reads sharing similar head base-pairs, a classic version of the Needleman-Wunsch algorithm in C++ (i.e., NW) with no matrix lines re-use but read deduplication, and EMBOSS NEEDLE [9], the most popular off-the-shelf Needleman-Wunsch algorithm implementation that does not deduplicate identical reads. The three aligners produced identical results.

Comparing the speed of each algorithm, we first verified that RecNW was on average 3.2 times faster than NEEDLE (range 2.4 − 5.8) on the limited real datasets (Table S1). In our extensive analysis with the realistic generated samples, we observed that across our 640 datasets RecNW was on average 4.4 times faster than NEEDLE (range 0.9–55.7; Paired t-test *p <* 10^−47^), and slower than NEEDLE for only two data sets out of 640 (Fig. S1). The speed of RecNW increased as the total number of reads and the amount of identical sequence in the reads increased, while the speed of NEEDLE stayed nearly constant (Fig. 1a, b, Fig. S1). The RecNW alignment speed of unique reads was highly correlated to the number of base-pairs reused (i.e., the number of lines the RecNW algorithm does not recompute) (*r*^2^ = 0.85, Fig. 1c, Fig. S2). This demonstrates how RecNW gains speed not only from simple deduplication of reads, but from the re-use of parts of the alignment matrices. This is a particularly useful feature for studies where mutations are restricted to a specific part of the sequence (Fig. S3), and the deduplication step provides greatest benefits for high read counts (Fig. S4).

**Figure 1:**
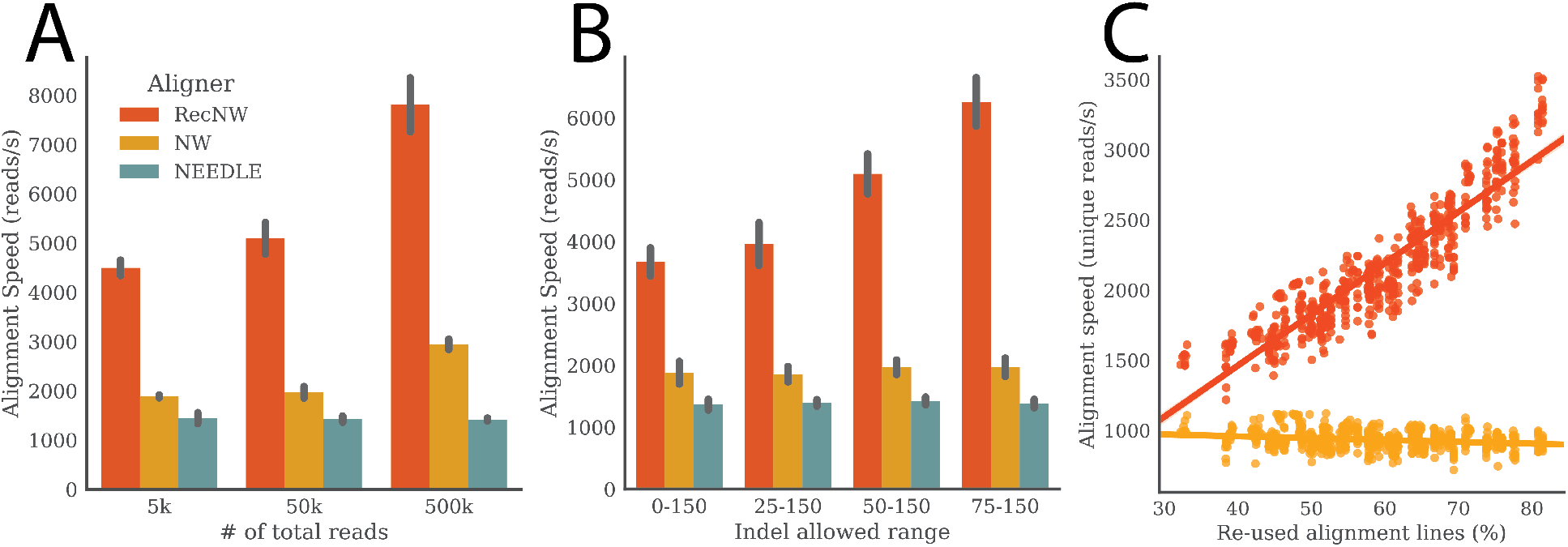
Alignment speed in total reads per second, between RecNW, NW deduplicating reads, and NEEDLE: A) as a function of the number of total reads for datasets with one indel per read restricted to positions in the 50−150 bp range; B) as function of the indel size allowed range for datasets of 50k reads with one indel per read. For instance, CRISPR experiments can yield large gaps at the cutting site. C) Alignment speed in unique reads per second as a function of the re-used alignment lines per dataset for RecNW and NW. This demonstrates how only RecNW has the feature to leverage the similarities between reads to increase the alignment speed.

In conclusion, RecNW provides identical alignment results to previous implementations of the Needleman-Wunsch alignment algorithm with substantially faster runtime especially for larger libraries of highly similar reads from specific loci. Thus, RecNW allows faster processing time for increasingly popular targeted deep sequencing applications without compromising accuracy.

## Acknowledgements

We would like to thank Ana Vasileva and Paul Hoffman for assistance with the FLCN data.

